# *Caledoniamyces theodoroui*, gen. nov., sp. nov., a bulbous anaerobic gut fungus from the bovine rumen

**DOI:** 10.64898/2026.10.15.759409

**Authors:** Susana Monino Ramos, Louisa Edge, Alice M. Buckner, Jessica L. Matthews, Stephane Kerangart, C. Jamie Newbold, Jolanda van Munster

**Author notes:** Corresponding author. School of Natural and Social Sciences, Scotland’s Rural College, King’s Buildings, West Mains Road, Edinburgh, EH9 3JG, UK. These authors contributed equally (Susana Monino Ramos, Louisa Edge).

## Abstract

Anaerobic gut fungi (AGF) are members of the microbiome of herbivore digestive tracts, where they are important degraders of plant material. Culture-independent diversity studies have indicated that a broad diversity of these fungi still remain uncultured so far. This limits understanding of their ecology, physiology and potential for biotechnological exploitation. Here we describe the isolation and characterisation of the first cultured representatives of the Neocallimastigomycota clade NY9 / YL1, previously proposed as genus based on culture-independent sequence diversity studies. Two fungal strains, RF6 and RF9, were isolated from Jersey cow rumen fluid, alongside representatives of other genera including *Orpinomyces* and *Neocallimastix*. Phylogenetic analysis of the RF6 and RF9 rDNA ITS1 region and D1–D2 domain of 28S LSU rDNA gene revealed formation of a well-supported genus-level cluster. This cluster also contained sequences deposited for NY9 and YL1 and was distinct from other described genera. Isolates RF6 and RF9 displayed small 1-2 mm colonies with a compact structure on solid medium. Microscopy showed bulbous rhizoidal cells encased in amorphous material, possibly extracellular polymeric substances, indicating that the genus represents the third known genus of AGF with bulbous morphology. Both endogenous and exogenous sporangia were observed, which produced mono-flagellated zoospores. Isolate RF6 was able to grow robustly using cellulose and wheat straw as sole carbon sources. For this new genus, for which the type strain is RF6, we propose the name *Caledoniamyces*, referring to the origin in Scotland, and the species name *C. theodoroui*.

**Highlights:**

- Isolation of a novel genus of anaerobic gut fungi (Neocallimastigomycota)
- First cultured representative of clade NY9/YL1
- The name *Caledoniamyces theodoroui* is proposed, with type strain RF6
- *C. theodoroui* has bulbous morphology and mono-flagellated zoospores

## 1. Introduction

Anaerobic gut fungi (AGF) are highly effective primary degraders of lignocellulose. They are commonly found within the gut of herbivores, where (together with protozoa, bacteria and archaea) they are part of a microbiome that enables conversion of plant material to available nutrients for their host (Gordon & Phillips, 1998; Krause et al., 2013). AGF are thought to degrade lignocellulose by leveraging both their penetrative hyphae for mechanical destruction and a wide range of carbohydrate active enzymes (CAZymes) for biochemical degradative activity. These CAZymes may be secreted as soluble enzymes, as known for aerobic fungi, or can be embedded in cellulosome-like multi-enzyme complexes, as reviewed by (Gilmore et al., 2015). AGF genomes encode the highest number of genes encoding plant-degrading CAZymes known in the fungal kingdom to date (Seppälä et al., 2017). This broad array of degradative tools may be instrumental in their effective lignocellulose degrading capacity, which is of vital importance to feed digestion, and holds potential to be exploited for sustainable green and white biotechnology.

Culture-independent studies and isolation efforts have indicated that AGF (phylum Neocallimastigomycota) are present in a broad range of herbivores (Kittelmann et al., 2012; Liggenstoffer et al., 2010; Meili et al., 2023, 2024), including in agriculturally important ruminants such as sheep and cattle (Davies et al., 1993; Lowe et al., 1987). In addition they can be found in pseudoruminants such as alpacas (Stabel et al., 2020) and hindgut fermenters such as horses, but also in tortoises (Pratt et al., 2024) and even ostriches (Vinzelj et al., 2025). Despite the isolation of many new AGF genera in the past decade (e.g. (Callaghan et al., 2015; Dagar et al., 2015; Hanafy et al., 2020; Joshi et al., 2018), a substantial diversity of AGF remains uncultured so far, with over sixty sequence-based candidate genera proposed (Meili et al., 2023). For one of these candidate genera, designated NY9 (Meili et al., 2023), at the same time recognised as sequence-based clade YL1 (Young et al., 2022), sequences were found usually in low abundance across samples from cattle, bison, lama and alpaca, yak, goat, buffalo. This suggests that that even for well-sampled agriculturally important ruminants, the representation of AGF genera in cultures remains incomplete.

The availability of cultured representatives of a broad range of AGF genera would enable studies aimed at gaining a deeper understanding into the ecology and evolution of these fungi. It would also enable functional and mechanistic studies to elucidate the fundaments of their biology, exploiting their functional diversity to accelerate development of applications. The presence of functional diversity within the AGF genera and species is suggested by specific fungal-host associations (Meili et al., 2023), as well as modulations of gut mycobiomes in response to distinct feed composition (Boots et al., 2013; Fliegerova et al., 2021). Functional studies have already highlighted that representatives of distinct AGF genera have different effects on plant material during its degradation (Shen et al., 2026), and distinct performances in improving biogas production (Vinzelj et al., 2020) alongside co-culture-based biobutanol and bio-butyrate production (Brown et al., 2022). Gaining further insight into the functional diversity of AGF thus provides promise for realising applications to advance biotechnology applications in addition to improving the feed efficiency of ruminants and sustainability of ruminant farming.

Here, we describe the isolation of the new genus and species *Caledoniamyces theodoroui* from the rumen fluid of Jersey cows. This genus corresponds to clades NY9/YL1, proposed based on culture-independent studies, demonstrating that previously uncultured genera can be isolated from cattle rumen fluid. The isolates were characterised by a bulbous rhizoidal morphology, which among Neocallimastigomycota previously has only been identified for the genera *Caecomyces* and *Cyllamyces*, and monoflagellated zoospores. The two isolates tested both grew robustly utilising a narrow range of substrates as their sole carbon source, including wheat straw and cellulose.

## 2. Methods

### 2.1 Sample collection and isolation

Rumen fluid was collected as described previously (Shen et al., 2026). Briefly, rumen fluid was manually collected from fistulated Jersey cows who where fed haylage twice daily, and immediately transported to the laboratory using a thermos flask. Animal procedures were carried out in accordance with the UK Home Office Animal (Scientific Procedures) Act 1986 and experimental protocols approved the University of Glasgow Animal Welfare and Ethical Review Board and the project license holder (project license PP3251909). During processing, the rumen fluid was strained through three layers of muslin to remove large particles.

Hungate tubes containing 6 mL medium C (based on (Theodorou et al., 2005), but modified to replace bactocasitone with tryptone), supplemented with 1% of wheat straw particles (milled to 0.5 mm diameter), penicillin (167 units mL^-1^) and streptomycin (167 µg mL^-1^, added as 100 µL Penicillin Streptomycin solution (Merck P4333) per tube) made under a CO_2_ atmosphere, were inoculated with 0.3 mL rumen fluid. Tubes were incubated stationary for 3 days at 39°C, after which 0.1 mL culture filtrate was used to inoculate roll tubes containing medium C with glucose and penicillin (167 units mL^-1^) and streptomycin (167 µg mL^-1^). After incubation for 3 days at 39°C, the tubes were inspected using a microscope to identify distinct fungal colonies, after which colonies were picked from the solid surface and transferred to liquid medium C supplemented with wheat straw (1%), penicillin (167 units mL^-1^) and streptomycin (167 µg mL^-1^). After growth, for selected isolates, colony picking via roll tubes was repeated to ensure isolation of pure fungal strains. Any remaining bacterial contamination was mitigated via addition of the following antibiotics sequentially in subsequent cultures: penicillin and streptomycin (Merck P4333, 100 µL per tube), kanamycin (80 µg mL^-1^), tetracycline (10 µg mL^-1^), moxifloxacin hydrochloride (25 µg mL^-1^), phosphomycin (330 µg mL^-1^), erythromycin (100 µg mL^-1^), ciprofloxacin (10 µg mL^-1^), vancomycin (100 µg mL^-1^), and - for isolate RF4 - chloramphenicol (50 µg mL^-1^). Obtained axenic fungal isolates were routinely maintained on medium C with wheat straw, supplemented with tetracycline (10 µg mL^-1^) (RF5, RF14, RF6, RF9) or chloramphenicol (50 µg mL^-1^) (isolate RF4). Cryogenic stocks were prepared by addition of 1.5 mL sterile anaerobic ethylene glycol stock solution, containing 49.7 g ethylene glycol, 155 mL clarified rumen fluid, 0.2 mL of resazurin solution (1 g L^-1^), 1.2 g NaHCO_3_ and 0.2 g L-cysteine-HCl, to two-day-old cultures, as described in detail for the CPeg protocol (Vinzelj et al., 2022).

### 2.2. Cloning and sequencing of genetic markers

Fungal biomass, derived from a 3-day old 6 mL culture with wheat straw, was used for DNA extraction. Biomass was retrieved from the culture by centrifugation at 15,000 *g* for 1 min, placed in 2 mL screw-cap tubes with 700 µL CTAB buffer, and cells were disrupted in a TissueLyser I (Qiagen) for 3 x 1 min at 50 Hz. Purification was performed following (Griffith et al., 2009) with the hot ethanol wash step replaced with ambient ethanol. Obtained DNA was used for subsequent LSU-based PCR-RFLP profiling, using PCR amplification of the 28S LSU rDNA region with NL1 and NL4 primers and digestion with AluI and Hpy188I, as described in detail in (Edge et al., 2026). For selected isolates, the internal transcribed spacer (ITS region) of the rDNA was amplified by PCR using the primers ITS1 and ITS4 as described by (Shen et al., 2026). ITS and LSU PCR products were cloned in pGem-T Easy vector (Promega) following manufacturer’s instructions, and nucleotide sequences were obtained via Sanger sequencing.

### 2.3. Phylogenetic analyses

Nucleotide sequences of D1-D2 28S LSU rDNA regions of AGF isolates were used to identify closest similar sequences using BLASTn against the core nucleotide database using default settings. Subsequently RF6 and RF9 LSU sequences were aligned with sequences from 28S rDNA (LSU) region of *Neocalllimastigales* clade YL1 (Young et al., 2022) and from NY9 (also designated NY09) (Meili et al., 2023), as well as representative sequences from cultured AGF genera, from type material or from the AF_LSU_database version 2.0 obtained from the Anaerobic Fungi Network (www.anaerobicfungi.org), using *Gonopodya prolifera* JN874506 as outgroup. Multiple sequence alignments were computed via Mafft v7.526 (Katoh & Standley, 2013), with alignment trimming using ClipKit v2.11.4 (Steenwyk et al., 2025) and maximum likelihood phylogenetic tree calculated using IQ-Tree v2.4.0 (Minh et al., 2020) using 5000 bootstraps. The pipeline was run using Phylogeny v2.0 (Dereeper et al., 2008), and visualised with MEGA 11, version 11.0.13 (Tamura et al., 2021). Distances between 28S LSU rDNA and ITS regions of AGF isolates were calculated using Blastn with a gap existence cost of 5; gap extension cost of 2; match score of 2; and mismatch score of −3 (Elshahed et al., 2022).

### 2.4. Amplicon sequencing

A 5 mL aliquot of rumen fluid was centrifuged for 10 min at 4500 rpm, at 4 °C. After removal of supernatant, the cell pellet was stored at -80°C until further processing. DNA was subsequently extracted as described by (Buckner et al., 2025) using the QIAamp Fast DNA Stool Mini Kit (Qiagen) combined with glass 0.1 mm PowerBead Tubes (Qiagen). DNA was used for amplicon sequencing with primers AGF-LSU-EnvS-F/R (Young et al., 2022). No amplification products were obtained for the negative control, and this was therefore omitted from further analysis. A minimum of 30k raw reads per sample was obtained via Illumina 250 bp paired-end sequencing. Bioinformatic analysis was performed as described (Edge et al., 2026) modified only to include an analysis step that trimmed the reference alignment used for initial contig alignment step to the amplified region using the pcr.seqs() command. Briefly, sequences were analysed using Mothur v.1.48.5 (Schloss et al., 2009). They were first assembled into contigs while removing primers and barcodes followed by quality trimming on sequences aligning to the amplified region of the Neocallimastigomycota reference alignment. Chimeras were then removed and the resulting operational taxonomic units (OTUs) classified using the AF_LSU_database(v2). The data were visualised via the R package phyloseq (McMurdie & Holmes, 2013).

### 2.5. Morphological characterization

Isolates RF6 and RF9 were grown in medium C with 10 g L^-1^ wheat straw or 5 g L^-1^ glucose as carbon source, for 2-7 days. Aliquots of cultures were retrieved, and mature fungal cells and zoospores were imaged using an Olympus BX53 light microscope equipped with a UPlanFL N 20x/50 ∞ /0.17/FN26.5 objective and a Canon EOS 60D camera. ImageJ (version 1.54g) (Schneider et al., 2012) was used for image processing and measuring cell dimensions.

### 2.6 Carbon source profiling

Isolate RF6 was pre-cultured for 3 days on medium C with wheat straw and tetracycline (10 µg mL^-1^), after which 0.3 mL culture was used to inoculate 6 mL medium C supplemented with a range of carbon sources. Glucose, mannose, arabinose, xylose, fructose, galacturonic acid, glucuronic acid, N-acetyl-glucosamine, fucose, sucrose, and maltose were all dissolved in dH_2_O at a concentration of 300 g L^-1^. Cellobiose and lactose were dissolved at 150 g L^-1^. Stock solutions were sterilised by filtration by passage through a 0.2 µm filter, and added to culture tubes containing sterile anaerobic medium to supply a carbon source at a final concentration of 5 g L^-1^. Polysaccharides were weighed out and added to culture tubes during medium preparation, to achieve 10 g L^-1^ for chitin (YC04085, Biosynth), inulin (from chicory, I2255, Merck), xylan (Corncob xylan, YC05381 Biosynth), galactomannan (from Guar, medium viscosity, P-GGMMV, Megazyme), glucomannan (from konjac, low viscosity, P-GLCML, Megazyme), and cellulose (microgranular, C6413, Merck). For pectin (Apple Pectin, 93854 Merck) and starch (potato starch, S4501 Merck) 5 g L^-1^ was used. Filter paper consisted of 60 mg strip of Whatman No. 1 filter paper. The wheat straw used was ground to a particle size of 0.5 mm. Cultures (n=6), with the exception of those grown with gluco- and galactomannan (n=4), were incubated for 4 days, stationary, in the dark at 39°C, after which growth was monitored via fermentation gas production, as standard practise in the field (Theodorou et al., 1995), using a Vernier LabQuest 2 data logger equipped with a pressure sensor, as well as by visual inspection. Cultures with fungal growth were sub-cultured twice to new ontaining the same carbon source, and their growth was monitored again after the third growth cycle. Cultures without growth at 4 days were monitored again after 6 days of growth to account for possible slow growth rates.

## 3. Results

### 3.1 Fungal isolations

Fungal enrichment cultures were performed with an inoculum of rumen fluid from fistulated Jersey cows, as preliminary amplicon sequencing analysis (Fig. S1) of the resident AGF community suggested that this rumen fluid contained a substantial amount of so far uncultured AGF genera, including the proposed genus NY9 (>45% of reads). Enrichment cultures in medium C were supplemented with wheat straw as carbon source and antibiotics penicillin and streptomycin to repress bacterial growth. Initial enrichment was followed by colony picking from roll tubes, which resulted in 10 AGF isolates. PCR-RFLP of the LSU region (Fig. S2), as we described recently (Edge et al., 2026), suggested that representatives of four distinct groups had been obtained: *Orpinomyces* sp. (isolate RF4), *Neocallimastix frontalis* (isolate RF12), a group of isolates with uncertain identity (RF5, RF7, RF11, RF14) described further elsewhere, and one group corresponding with proposed genus NY9 (RF6, RF9, RF10, and RF13). Based on this initial assessment, selected isolates were purified to axenic culture and further characterised. Subsequent cloning and Sanger sequencing of LSU regions confirmed the identity of isolate RF4 as *Orpinomyces joyonii* (99.74-100% identity to *O. joyonii* B1 Genbank PX908539.1). Notably, initially obtained sequences belonging to isolates RF6 and RF9 had high similarity (98-99%) to sequences previously deposited for proposed genus NY9 (Meili et al., 2023), also referred to as YL1 (Young et al., 2022), suggesting these isolates may be the first isolated representatives of this genus.

### 3.2 Phylogenetic analysis

Additional clones of the D1-D2 28S (LSU) rDNA regions, as well as ITS regions were obtained for RF6 and RF9. Sanger sequencing of LSU region clones for RF6 indicated intra-isolate variation of 0-0.26% (n=14) and 0-0.13% for RF9 (n=5), and 0-0.25% intra-isolate variation.

BLASTn analysis of obtained LSU sequences indicated high similarity to sequences deposited for proposed genus NY9, with 98.7-96.6% identity to OP253940-OP253955 and OP253753-OP25375356. Comparison with sequences from *Caeocomyces communis* var. *churrovis*, identified via BLAST as the nearest closest cultured species, showed up to 95.4% sequence identity. Following guidelines for sequence divergence of at minimum 3% for the closest described taxa (Elshahed et al., 2022), this suggests the RF6 and RF9 isolates are the first cultured representatives of the genus previously designated as NY9.

Phylogenetic analysis (Fig. 1). confirms LSU sequences of isolates RF6 and RF9 cluster together with sequences obtained in culture-independent studies for proposed genera/clade NY9 and YL1. The sequences together form a well-supported clade (bootstrap value of 100), distinct from other genera and established AGF families, of which the closest family is

**Fig. 1.**
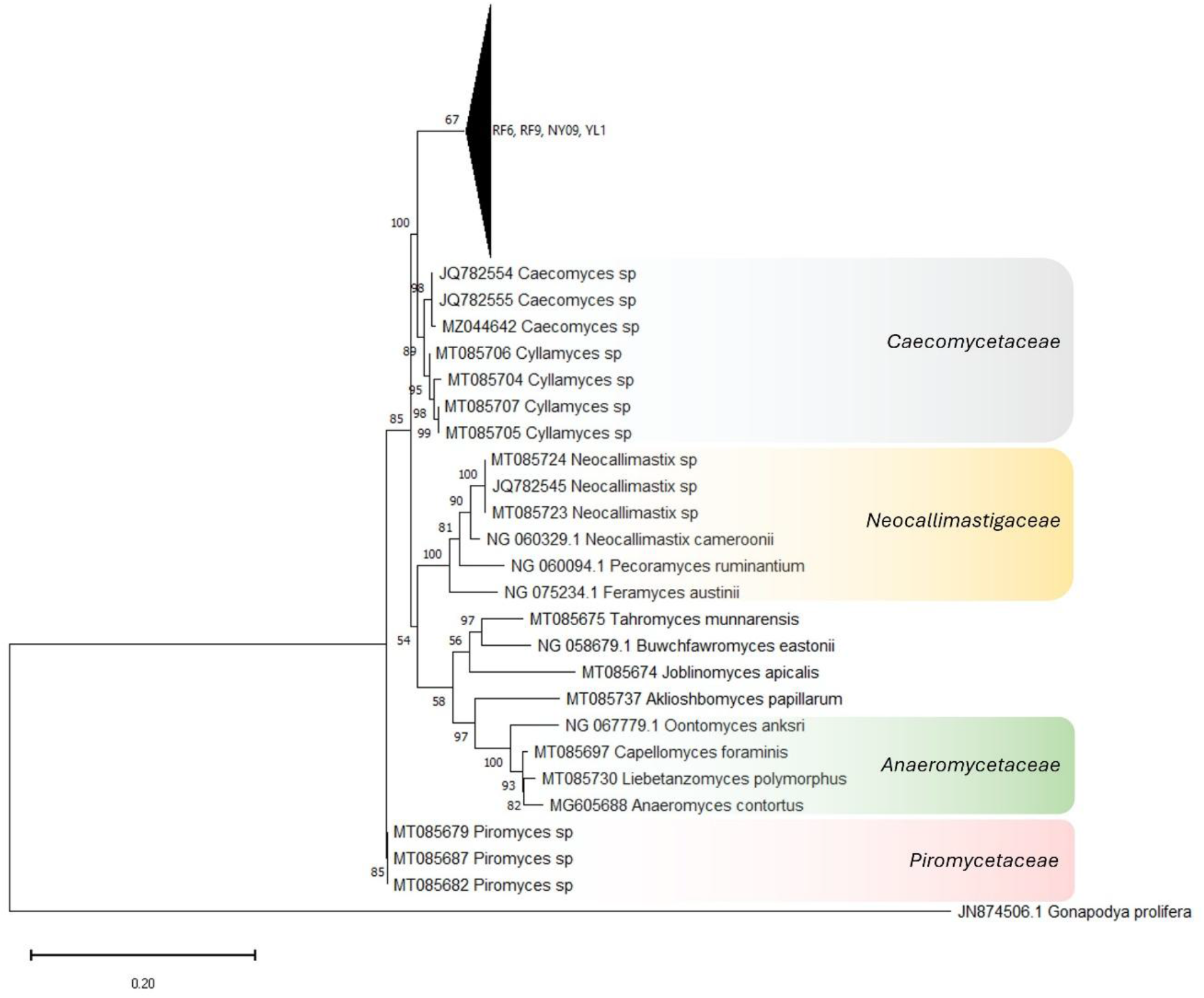
Maximum Likelihood tree of Neocallimastigomycota based on D1-D2 28S rDNA sequences. Numerical values report bootstrap support, recognised families are indicated in coloured bars, the triangle groups sequences from isolates RF6, RF9 and those obtained from culture independent studies describing clades NY09 (GenBank accessions OP253753-OP253756 and OP253940-OP253955) and YL1 (GenBank ON758305-ON758307, ON682552-ON682557, ON831951-ON831956, ON819093, ON819099, ON819132, ON819144, ON819145, ON819159, ON819163), an expanded tree is provided as Fig. S3.

*Caecomycetaceae*. We examined the relation between sequences in the clade (Fig. S2) as (Young et al., 2022) indicated clade YL1 may include several species, and found a sequence divergence of up to 2.8% across the clade, providing scope for species delineation. However, bootstrap support for separation into groups consistent with species-level sequence divergence was limited (69% or lower). When more sequences, for example from additional isolates become available, this should be re-assessed.

Sanger sequencing of cloned ITS sequences from isolates RF6 and RF9, and subsequent BLASTn analysis indicated the intra-isolate sequence divergence was 0-0.41% (n=7). Closest sequences from cultured representatives were from *Caecomyces communis* var. *churrovis*, at 92.9% identity, well over the 5% ITS sequence divergence threshold recommended for novel AGF genera (Elshahed et al., 2022) .

We thus propose to accommodate the isolates RF6 and RF9 in a novel genus, *Caledoniamyces theodoroui*, with the genus name referring to Caledonia, the ancient Latin name for Scotland, where the isolates were obtained, and the species epithet *theodoroui* honouring microbiologist and anaerobic fermentation expert Prof. Mike Theodorou.

### 3.3 C.theodoroui represents the third genus with bulbous morphology

The colonies of RF6 and RF9 grown on solid medium in roll tubes were small, initially below 2mm diameter, but later with cells dispersing outwards. Colonies had an irregular, somewhat circular shaped circumference, without visible filamentous hyphae (Fig. 2A). Visual inspection via microscopy showed colonies consist of bulbous structures, similar to those of *C. communis* (Fig. 2B).

**Fig. 2.**
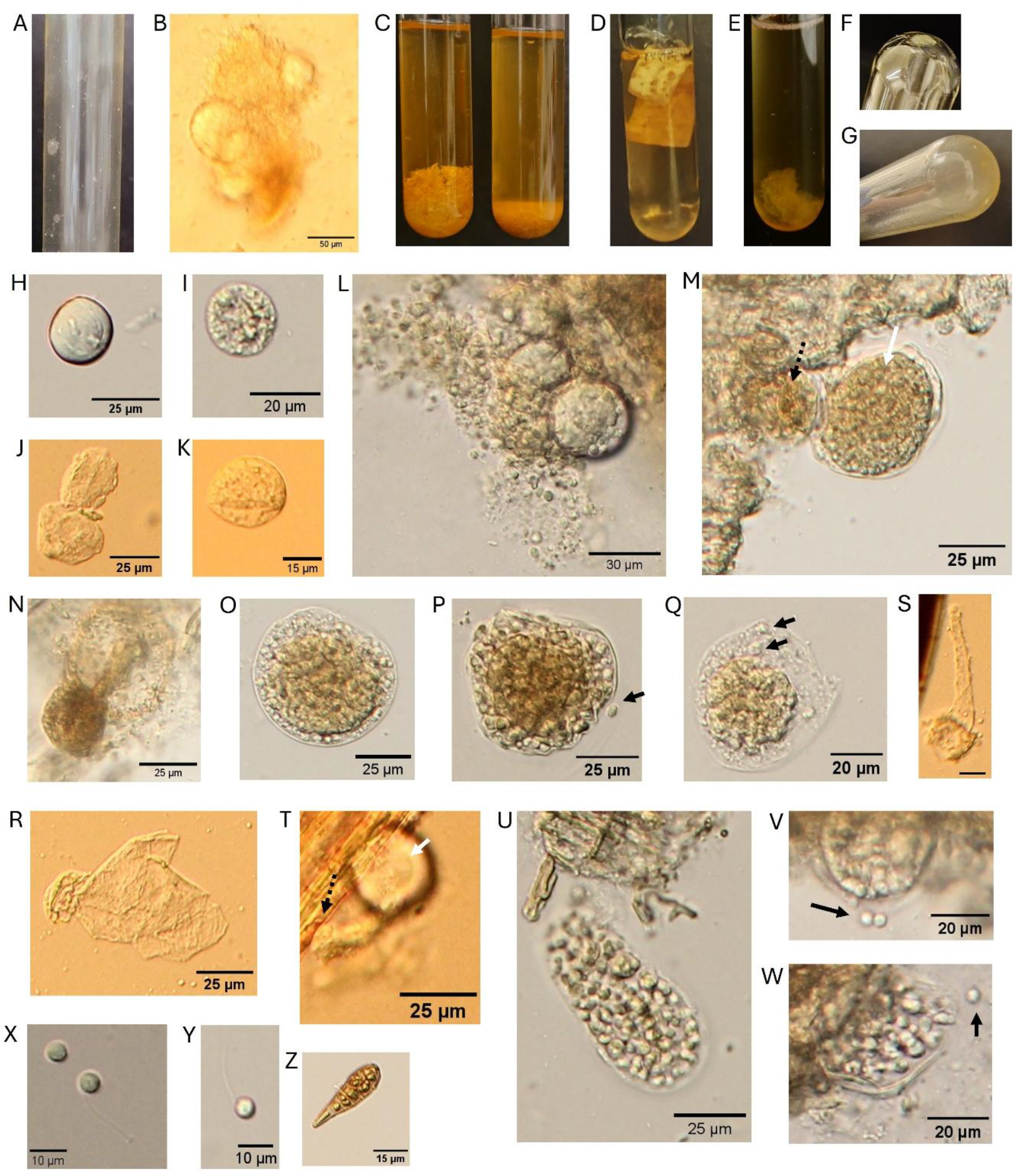
Morphological descriptions of *Caledoniamyces theodoroui*, isolate RF6 unless indicated otherwise. A,B colonies in 3d old roll tubes; C cultures in liquid medium, 3d old, wheat straw as carbon source (left with RF6, right abiotic medium); D filter paper as carbon source, 6d old; E RF9, glucose as carbon source, 3d; F,G on lactose as carbon source, 5d and 9d respectively, biofilm on glass with medium decanted; H-K bulbous rhizoids or thalli, H,K 3d on glucose, I,J 7d on wheat straw; L 3d old, glucose, unstructured material layer visible on cells; M-U 7d on wheat straw, M bulbous rhizoidal foothold cell (black arrow) with sporangium (white arrow); N bulbous rhizoid with endogenous sporangium; O-Q sporangia, zoospores visible inside or after release (black arrows); R bulbous rhizoidal cell with remnants of endogenous sporangium after zoospore dispersal; S rhizoidal cell with tubular, slightly tapering sporangiophore, scalebar 15 µm; T rhizoidal cell attached to straw (black dashed arrow) with sporangiophore and exogenous sporangium (white arrow); U sporangium filled with zoospores; V-Y 3d old cultures on glucose, V,W sporangium releasing zoospores (black arrow) through apical pore; X,Y mono-flagellated zoospores; Z, possible resting stage or spore, 7d culture on wheat straw.

During growth in liquid medium with wheat straw as carbon source (Fig. 2C), RF6 loosely assembled straw particles together, forming clumps of straw and mycelium that were easily dispersed, especially compared to biomass rafts that were formed by *Neocallimastix frontalis, Feramyces austinii*, and other filamentous AGF. On filter paper, colonies were visible as more transparent spots, with the paper curling up in later growth stages (Fig. 2D). Gross morphology of cells in media with soluble carbon sources (Fig. 2E, glucose) was initially grainy and amorphous, with biomass yellowish to light brown in colour. Fungal mycelium adhered strongly to the glass tube surface forming a sticky layer (Fig. 2F, G), that dissociated from the tube when grown into a thick layer of biomass (Fig. 2E).

Microscopy demonstrated that rhizoidal cells were bulbousfilamentous hyphaea were not observed. The cells were spherical (Fig. 2H, I) to elliptical (Fig. 2K) and 16-27 µm in diameter when observed alone or as part of a small cluster of two (Fig. 2J) or three cells. Cells displayed both smooth and structured surfaces. Most of the rhizoidal cells, however, were observed as part of larger clusters covered in a non-structured compound (Fig. 2L). This compound could potentially be an extracellular polymeric substance that may facilitate the observed strong biofilm formation.

Endogenous and exogenous sporangia were observed, with a single sporangia per rhizoidal foothold. The endogenous sporangia (Fig. 2N-R), 39-65 µm in diameter, were regularly observed as being separated from their bulbous foothold. They ranged from almost globose cells with an even dark centre (Fig. 2N,O) to ovoid or turbinate with dark structured centre with visible zoospores (Fig. 2P,Q). Exogenous sporangia (Fig. 2S-U), were obpyriform or ellipsoid in shape and up to 38 µm wide and 71 µm length, and found at the end of tubular sporangiophores of up to 63 µm length (Fig. 2S, T). Zoospores, which were released from apical pores or openings in the sporangia (Fig. 2V,W), were mono-flagellated and often spherical in shape, though irregular and elliptical shapes were observed. The average zoospore diameter was 4.8 ± 0.6 µm (n=50)and average flagellum length was 17.4 ± 1.6 µm (n=25).

### 3.3 Carbon source usage

Isolate RF6 was able to grow vigorously on glucose, fructose, and xylose, however it was unable to use any of the other tested monosaccharides as sole carbon source (Table 1). Growth assays on dissacharides showed that cellobiose supported biomass and gas production levels equivalent to monossacharides whereas lactose utilisation proceeded at a slower rate, with full biomass production observed after > 1 week (Fig. 2G). Cellulose was the only polysaccharide RF6 was able to use as sole carbon source, both when supplied as particles or as the more complex three-dimensional material filter paper. Collectivelly, our results suggest a limited range of carbohydrates can be utilised by isolate RF6 as sole carbon source.

**Table 1.** Growth of isolate RF6 on medium C with a range of carbon sources.

| Substrate type | No growth | Growth |
| --- | --- | --- |
| Monosaccharide | Mannose | Glucose |
|  | Fucose | Fructose |
|  | Arabinose | Xylose |
|  | N-acetyl-glucosamine |  |
|  | Glucuronic acid |  |
|  | Galacturonic acid |  |
| Disaccharide | Maltose | Lactose |
|  | Sucrose | Cellobiose |
| Polysaccharide | Xylan | Cellulose |
|  | Pectin |  |
|  | Starch |  |
|  | Galactomannan |  |
|  | Glucomannan |  |
| Complex |  | Wheat straw |
|  |  | Filter paper |

### 3.6 Taxonomy

*Caledoniamyces* Susana Monino Ramos, Louisa Edge, Alice Buckner, Jessica Matthews, Stephane Kerangart, Jamie Newbold, Jolanda van Munster, gen. nov. Mycobank ID: 865760

Typification: *Caledoniamyces theodoroui* Susana Monino Ramos, Louisa Edge, Alice Buckner, Jessica Matthews, Stephane Kerangart, Jamie Newbold, Jolanda van Munster (holotype)

Ethymology: *Caledonia* is ancient Latin for Scotland, where the fungus was isolated; *myces* is Greek for fungus.

Obligate anaerobic fungus that grows via bulbous thalli. Produces endogenous sporangia as well as exogenous sporangia on tubular sporangiophores, which release mono-flagellated zoospores. The sequences QB178310-QB178316 and QB178293-QB178309 represent the clade’s ITS1-5.8S rDNA-ITS2 and D1/D2 28S rDNA regions. The genera most similar genetically are *Cyllamyces* and *Caecomyces*, which both also produce bulbous thalli, as reviewed by (Hanafy et al., 2021).

*Caledoniamyces theodoroui* Susana Monino Ramos, Louisa Edge, Jessica Matthews, Alice Buckner, Stephane Kerangart, Jamie Newbold, Jolanda van Munster (holotype) Mycobank ID: 865762

Typification: The holotype shown in Fig 2H-Z, is from a 7-day-old culture of strain RF6 grown in medium C with wheat straw, the isolate obtained from rumen fluid from a fistulated Jersey cow from the west of Scotland, in March 2026 in Edinburgh, Scotland (55.922, - 3.177). Ex-type strain: RF6. GenBank QB178310, QB178311, QB178313-QB178315 (ITS1-5.8S-ITS2) and QB178293-QB178305 (D1/D2 28S). The type culture is preserved cryogenically as well as in continuous culture at Scotland’s Rural College (SRUC) in Edinburgh.

Ethymology: The species epithet *theodoroui* honours refers to Dr Mike Theodorou, an expert in anaerobic gut fungi with an extensive track record of leading development of anaerobic fungi in biotechnology applications.

An anaerobic fungus, forming compact irregular shaped colonies on solid media in roll tubes. Liquid growth is as biofilm with bulbous rhizoidal cells encased in amorphous material, firmly attached to surfaces. Zoospores are spherical, 4.8 ± 0.6 µm in diameter, monoflagellated, with flagella 17.4 ± 1.6 µm long. Rhizoidal cells are bulbous, elliptical or spherical, 16-27 µm in diameter.Produces endogenous sporangia, 39-65 µm in diameter, with large dark fusiform centres and exogenous sporangia of up to 71 µm in length on tubular sporangiophores.

Additional specimens examined: Isolate RF9 (GenBank QB178312, QB178316 and QB178306-QB178309), isolated from the same rumen fluid sample.

## 4. Discussion

Here we proposed to accommodate the AGF isolates RF6 and RF9 in a novel genus and species, *Caledoniamyces theodoroui*. Based on sequences of the 28S rDNA region, which is recognised as effective phylogenetic marker for Neocallimastigomycota (Hanafy et al., 2023; Young et al., 2026), the isolates form a clade supported by stringent bootstrapping values, distinct from so far recognised genera and families. Furthermore, sequence divergence of the *C. theodoroui* isolates D1-D2 28 sDNA regions to the nearest described genus was 4.6%, confirming the isolates’ placement in a separate genus. While most AGF genera harbour fungi that have filamentous rhizoids, *C. theodoroui* had bulbous rhizoidal cells, as described only for the AGF genera *Caecomyces* (Gold et al., 1988) and *Cyllamyces* (Ozkose et al., 2001). However, *C. theodoroui* isolates had clear morphological features that distinguish them from *Caeocmyces* and *Cyllamyces* representatives. Compared to the rhizoidal foothold, *C. theodoroui* had singular large exogenous sporangia, not dissimilar as those seen for *Neocallimastix frontalis* or *Piromyces communis*, as well as larger endogenous sporangia, while *Caecomyces* and *Cyllamyces* sporangia were comparatively smaller (Hanafy et al., 2021; Ozkose et al., 2001; Wubah et al., 1991). While *Cyllamyces* thalli are polycentric, with multiple sporangia per rhizoidal foothold, and nuclei migrating in both rhizoidal and sporangial cells, *Caecomyces* thalli are monocentric (Hanafy et al., 2021; Ozkose et al., 2001; Wubah et al., 1991). For *Caledoniamyces*, only one sporangium per cell was observed, suggestive of *Caledoniamyces’* monocentric development. However, the location of nuclei could not unambiguously be determined for *Caledoniamyces*. Taken together, fungal isolates RF6 and RF9 are thus representatives of both a morphologically and phylogenetically distinct genus.

Gross morphology of *C. theodoroui* thalli in cultures resembled that of filamentous genera. Despite the absence of hyphae, cells clustered together in a biofilm, whereas other bulbous species are described as forming small sandy grain-like structures. (Hanafy et al., 2021). Although confirmation of identity would be required, it is tempting to speculate that the unstructured material found on the *C. theodoroui* cells may be extracellular polymeric material (EPS), a material that is involved in biofilm formation in a range of fungi (Beauvais et al., 2007; Flemming & Wingender, 2010). This EPS may be responsible for the observed biofilm-like growth in liquid culture and in the rumen environment could allow the fungus to attach firmly to feed materials ingested by the animal host.

The genus *Caledoniamyces* incorporates sequences of candidate genera NY9 (Meili et al., 2023), and clade YL1 (Young et al., 2022), which offers valuable insights into the distribution and prevalence of the genus. In these culture independent studies, NY9 sequences were found relatively abundant (>10% of sequences) in faecal samples of a goat, yaks and cows, and in lower abundances (3-10%) in samples from buffalo, camel, horse, oryx, and sheep (Meili et al., 2023). However, it should be noted that sequences from the genus were detected only in a fraction of the replicate samples from these host species. In faecal samples from camel species, NY9 sequences were reported ranging from 0-30% relative abundance (De Silva et al., 2026). As clade YL1 was found with 4-6% abundance in cow, bison, lama and alpaca faecal samples (Young et al., 2022) together this covers mainly, but not exclusively, samples from foregut fermenters of family *Bovidae* and pseudoruminants of family *Camelidae*. We detected abundant *Caledoniamyces* sequences in, and isolated the fungus from, rumen fluid of a cow, matching the previously reported hosts. The varying prevalence of *Caledoniamyces* in host animals reported by (De Silva et al., 2026; Meili et al., 2023; Young et al., 2022) may be explained by factors known to influence gut AGF community composition, such as animal host diet (Boots et al., 2013; Fliegerova et al., 2021). *C. theodoroui* was able to use only a small selection of carbohydrates as a sole carbon source, including glucose, fructose, xylose, the polymer cellulose, and its breakdown product cellobiose. This substrate scope suggests some specialisation in sole carbon source use, not dissimilar as found for *Caecomyces* and *Neocallimastix* (Matthews et al., 2026), but in contrast to representatives of other AGF genera with broad substrate usage such as *Feramyces* (Hanafy et al., 2018). How such differences in carbon source specialisation affect fungal niche and functions in the rumen microbiome or their cross-kingdom interactions is currently underexplored (Shen et al., 2026). However, with cultured isolates of a growing number if AGF genera in hand, we can start to explore such functional differences. This provides fundamental insights that will contribute to the use of AGF’s unique capabilities and characteristics to advance applications in renewables-based biotechnology or ruminant nutrition.

## CRediT authorship contribution statement

**SMR**: conceptualisation, methodology, validation, formal analysis, investigation, data curation, writing-review & editing, visualisation; **LE**: methodology, formal analysis, investigation, data curation, writing-review & editing; **AMB**: conceptualisation, methodology, validation, formal analysis, investigation, data curation, writing-review & editing, visualisation; **JLM**: investigation, data curation, writing-review & editing; **SK**: investigation, **JN**: funding acquisition, conceptualisation, writing-review & editing. **JvM**: conceptualisation, methodology, validation, formal analysis, investigation, data curation, writing - original draft, writing-review & editing, visualisation, supervision, project administration, funding acquisition.

## Data availability

Sequences of D1-D2 28S rDNA region of *Orpinomyces joyonii* isolate RF4 are available via GenBank accession numbers QB156045-QB156047, sequences of D1-D2 28S rDNA region of *Caledoniamyces theodoroui* isolate RF6 as GenBank QB178293-QB178305 and for isolate RF9 as GenBank QB178306-QB178309. Sequences for clones of ITS1-5.8S-ITS2 region of *Caledoniamyces theodoroui* isolate RF6 are available as GenBank QB178310, QB178311, QB178313-QB178315 and for isolate RF9 as GenBank QB178313, QB178316. The fungal genus and species holotype are available as Mycobank accession number [865760, 865762].

## Conflict of interest statement

Declarations of interest: none

## Acknowledgements

The authors thank farm staff for sample collection. This work has been supported by funding from the Royal Society via URF\R1\231686, the UKRI Biotechnology and Biological Sciences Research Council (BBSRC) through EASTBIO DTP, grant number BB/T00875X/1.

## Appendix A. Supplementary data

Supplementary data is available as pdf associated with this article.

## Notes

### Competing Interest Statement

The authors have declared no competing interest.

