## Supplementary information for "*Caledoniamyces theodoroui*, gen. nov., sp. nov., a bulbous anaerobic gut fungus from the bovine rumen"

Supplementary data

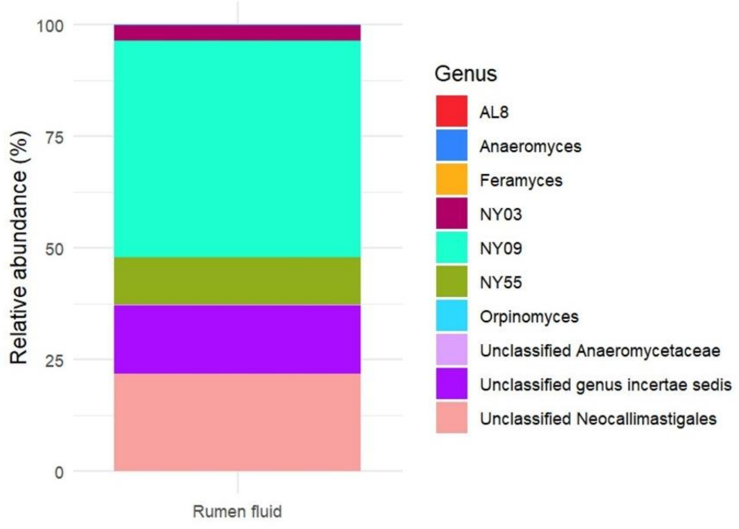


**Fig. S1.** Culture independent analysis of AGF community in a rumen fluid sample from jersey cows, as used for fungal isolations. Classification of obtained 33 OTUs to genus level is displayed, indicating an abundance of so-far uncultured AGF including the NY09 genus-level clade.

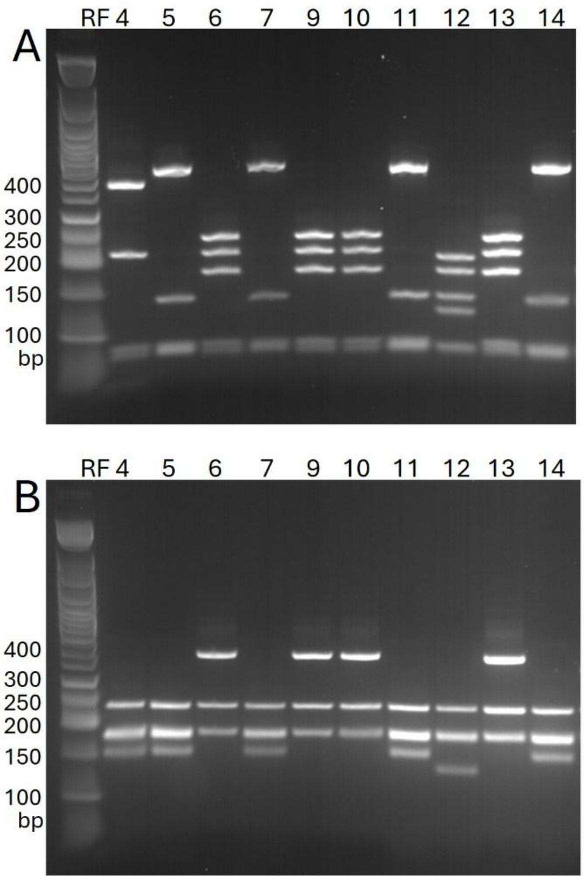


**Fig. S2.** PCR-RFLP based on LSU region of initially obtained fungal isolates from rumen fluid. Isolates are indicated as RF4-RF14, size of selected bands of the 50 bp DNA ladder is indicated in base pairs. A, RFLP pattern after digestion with AluI; B, digestion with Hpy188I.

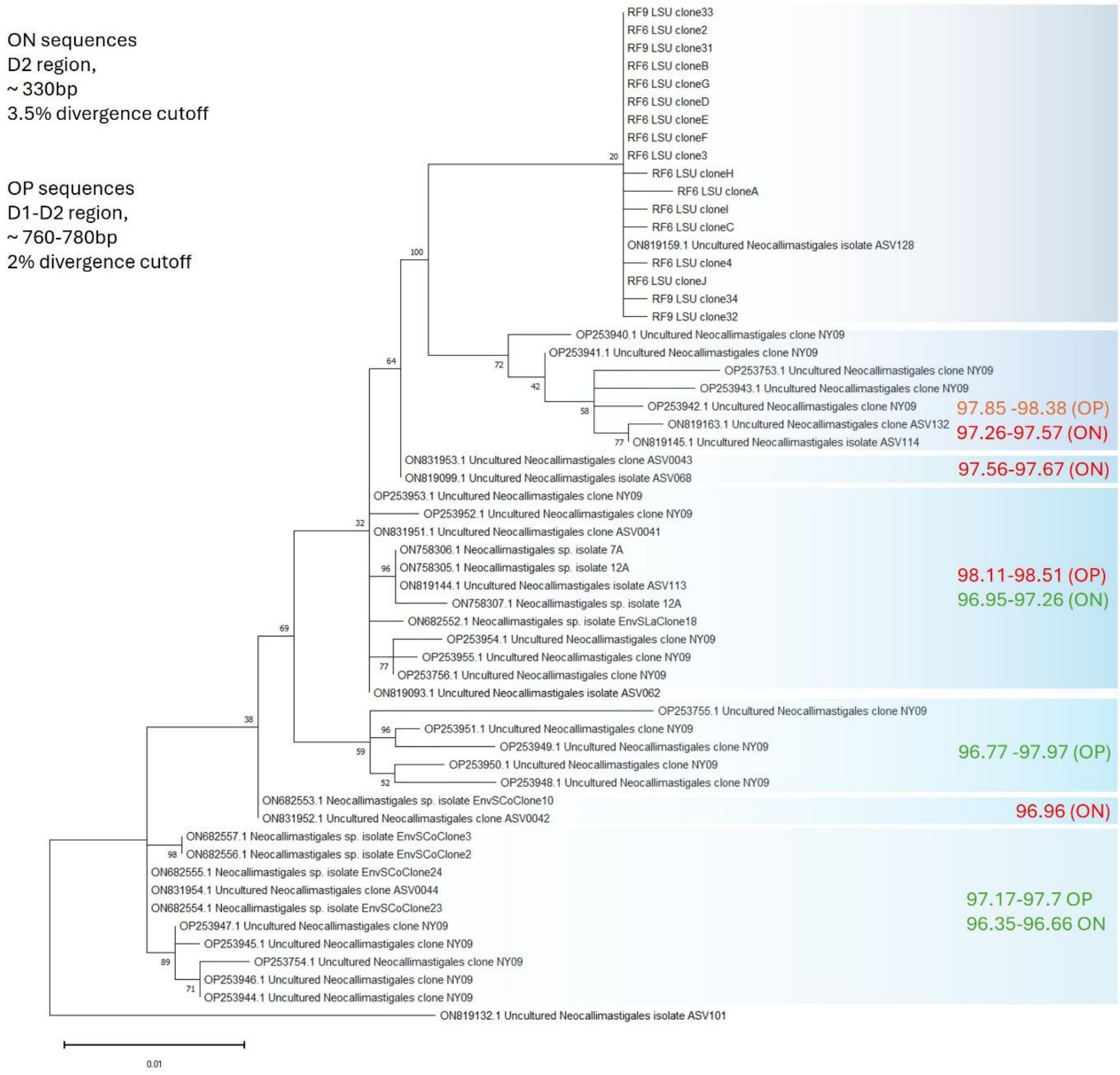


**Fig. S3**. Maximum Likelihood subtree, based on D1-D2 28S rDNA sequences and derived from Fig. 1, displaying only the expanded genus-level clade containing sequences from isolate RF6, RF9, and previously proposed NY09 and YL1 clade sequences. Numerical values on the tree report bootstrap support, green and red labelling indicates % sequence identity in the labelled subclade as compared to RF6 and RF9 isolate sequences. Green indicates below, and red above sequence divergence cut-off proposed by Meili *et al.* (2023) as species level separation.
